# MUTALISM BETWEEN *Ambystoma maculatum* AND *Oophila amblystomatis* IS DEPENDENT ON LIGHT INTENISTY DURING PHOTOSYNTHETIC GROWTH

**DOI:** 10.64898/2026.09.27.754580

**Authors:** Laura Palumbo, Glenn J. Tattersall, Wylan Clitheroe, Patrick Moldowan, Harvey Bates, Alonso Zavefer

## Abstract

The most studied symbiosis between land vertebrates and algae, is the one between *Ambystoma maculatum* and *Oophila amblystomatis*, where the latter colonizes the amphibian’s eggs. Although clear benefit given by algal growth for the salamander embryo has been observed, mutualism is not widely accepted. Mutualistic hypotheses posit benefits from the host to the algae which include protection, N- and C-sources, but no experimental evidence has been obtained. In fact, recent studies suggest that the algae are stressed during this relationship and propose fermentation rather than photosynthesis as main driver for algal growth. In this work, we analyzed embryonic host developmental rate, algal and embryo growth and photosynthetic function of eggs housed at different light intensities. We provide evidence that photosynthetic growth is the main driver for Oophila and that higher algal growth yields faster developmental rate and embryo size. Also, we provide strong evidence that rates of photosynthetic electron transport (and the concomitant O_2_ evolution) must be higher than those of consumption by the embryo for this symbiosis to be mutualistic. Since photosynthetic rate is dependent on light, this work illustrates the importance of light intensity in this symbiotic relationship.

## 1. Introduction

Symbiotic relationships between eukaryotic algae and animals are well documented, although there are few examples of endosymbiosis of microalgae and land-dwelling amphibians Yang et al. (2022). All well documented cases involve amphibians establishing symbiotic associations with facultative endosymbionts of the taxa Oophila (order Chlamydomonales)(Yang et al. 2022), specifically during the early embryonic stages of development (Bishop and Miller 2014). Among all known examples, the association between *Oophila amblystomatis* (syn. *Chlorococcum blastomata*) and *Ambystoma maculatum* is the most extensively studied, with over 80 years of reports (Bishop and Miller 2014). Despite this long-standing recognition and the uniqueness of the phenomenon, our understanding of this symbiosis remains limited (Yang et al. 2022). In this interaction, green algae colonize the interior of the egg capsule (Kerney et al. 2011). It has also been demonstrated that some algal cells invade the embryo’s body (Kerney et al. 2011), forming intracellular cysts (Kerney et al. 2011). The presence of Oophila DNA in the oviduct of adult females suggests that vertical transmission of the algae from mother to offspring may be possible (Kerney et al. 2011).

A conventional view defines this symbiosis as mutualism between the amphibian and autotrophic microbes, where net benefits for both parties exist, and herein referred as the *mutualism paradigm* (Hutchison and Hammen 1958; Yang et al. 2022). This paradigm has been explained by multiple hypothesises, *e.g.*, protection of the algae by the host, CO_2_ and/or Nitrogen sources from the host to the algae (Small et al. 2014), while the autotrophs may provide the host with sugars, and O_2_ provision and de-nitrification by the algae (Small et al. 2014). Since the 1940’s (Gilbert 1942, 1944; Hutchison and Hammen 1958), multiple experiments subjecting the holobiont (host organism and associated microbes) to light and dark treatments have demonstrated a clear benefit for the salamander host in terms of development and survival (Hutchison and Hammen 1958; Tattersall and Spiegelaar 2008; Small et al. 2014). However, evidence supporting a clear benefit for the algae is much more ambiguous (Burns et al. 2017), and much less attention has been given to the physiology of the algae.

An alternative view questions the net benefit for Oophila, suggesting that the algae may, in fact, experience physiological stress due to the internal conditions of the egg. Given the hypoxic environment that egg masses often encounter underwater, it was initially hypothesized that Oophila shifts its bioenergetic metabolism from oxidative phosphorylation to fermentation (Burns et al. 2017), a common response of clade Chlamydomonadales. This hypothesis was later supported by transcriptomic analyses showing that genes associated with fermentation and stress response are expressed by the algae during the salamander’s development (Burns et al. 2017). This alternative view does not negate benefits for the host but proposes no net bioenergetic benefit for Oophila from this association, here in referred as the *stress paradigm*. If the stress view is correct, a logical conclusion is the redefinition the symbiosis is between Oophila-spotted salamander as amensalistic. Moreover, recent studies using stable isotope tracing have revealed that the salamander embryo and the alga compete for available inorganic carbon within the egg capsule, with the embryo’s own carbon fixation processes potentially masking any minor transfer of photosynthetically derived carbon from the algae (Burns et al. 2020).

Interpretation of the evidence supporting the stress paradigm contradicts photobiological findings supporting mutualism as the main driver of this symbiosis. Studies of the photophysiology of Oophila during the first 25 days of development have shown that Photosystem II (PSII), is active in the algae during the early stages of development. In addition, O_2_ produced photosynthetically by the algae is utilized by embryos of the salamander covering a portion of the O_2_ demand (Gilbert 1944; Bachmann et al. 1986; Pinder and Friet 1994). However, if algae would be growing via fermentation, a complete cessation of the photosynthetic reactions would be required as demonstrated in *Chlamydomonas reihartii* (Gfeller and Gibbs 1984), the best studied Clamidomonales species.

If Oophila growth is a light dependent process (Small et al. 2014; Olivier and Moon 2010), different light environments should have an impact on the algae, affecting the host development as well. Indeed, light treatment experiments, where the holobionts are subjected to different photoperiod, have demonstrated impact on this symbiosis (Tattersall and Spiegelaar 2008; Small et al. 2014; Small and Bishop 2020). For example, holobionts exposed to a photoperiod of 24 h accelerated hatching for 30 days when compared to those reared under a dark and 12:12 h photoperiods in experiments done at 10°C(Tattersall and Spiegelaar 2008). In another study, dark treatment holobionts showed slower developmental responses when compared to a 14:10 h and 24 h photoperiod (Small and Bishop 2020). In addition, in this study it was observed that chlorophyll content was undetectable in the dark treated samples (Small et al. 2014; Small and Bishop 2020). Explaining these results under the stress paradigm is not possible, as Chlamydomonales synthesize chlorophyll in the dark under anaerobic growth conditions (Duanmu et al. 2013). Therefore, holobionts kept in the dark should display some level of chlorophyll phenotypically if the stress paradigm is correct.

Reconciling the evidence supporting both paradigms would require further investigation, particularly as most of the work supporting the mutualistic paradigm focuses on the salamander physiology. For this reason, in this work the photophysiology of the symbiont is studied *in vivo*, *in situ,* and in a non-invasive manner. We monitored photosynthetic activity of the light reactions, in particular PSII activity and the photosynthetic electron transport rate (ETR) in the context of the rate of development of the symbiont. Then, the obtained experimental data is contrasted against the literature where we model different scenarios of O_2_ evolution (under our experimental conditions) providing an alternative view to explain photoheterotrophic growth under the mutualistic paradigm.

## 2. Methods

### 2.1 Biological Material

Fifteen freshly laid egg masses of the spotted salamander (*Ambystoma maculatum*) were collected between May 4–6, 2025, from Bat Lake in Algonquin Park (45.576927° N, –78.522514° W; Nipissing District, Northeastern Ontario, Canada). Daily observations during the week prior to collection indicated that the eggs had been laid less than 24 hours before sampling.

Following collection, the egg masses were transferred indoors to a cabin maintained at ambient outdoor temperature. They were placed in plastic vessels filled with Bat Lake water, under room light and aerated with bubblers for three days until our team was able to transfer them to our laboratory. At our lab, > 10 eggs were separated from each of the 14 egg masses, while one mass was left untouched as a control. Because each egg represents a single mother, to minimise bias from maternal effects, eggs were subsequently arranged into four groups that later would be subject to different light doses. LEDs were positioned at a height far enough from the eggs to avoid thermal effects by their circuitry. Temperature effects were assessed using a thermocouple probe placed at the same distance from the light source as the eggs. Measurements from this probe were compared with readings from additional probes positioned in other areas of the temperature-controlled room.

Each experimental group therefore had eggs from five different mothers (4 eggs of the same egg mass), randomly selected, and started with a total of 20 eggs. Some eggs were ultimately unviable and were removed from the study and kept away from the experimental eggs to avoid fungal propagation. It is worth noting that Oophila continue to grow despite the presence of the fungus (Figure S1).

Individual eggs were transferred to cylindrical glass vials compatible with a Walz Water PAM fluorometer for use in the experimental design. Both individual eggs and intact egg masses were maintained in a temperature-controlled room at 15°C and illuminated with broad-spectrum LEDs (Tabletop Mini Garden 30.07 PPF LED, Utilitech, China). Remaining egg masses were housed separately in autoclaved pipet tip boxes (Fisher Scientific, USA) filled with Bat Lake water, which was changed weekly and were left untouched as controls to confirm embryos developed satisfactorily.

Once larvae hatched, individuals were euthanized using buffered MS-222. Experimental use of salamander eggs was approved by the Brock University Animal Care Committee (AUP: 25-03-01).

### 2.2. Experimental Design

Three egg groups (N=20 per group) were exposed to a photoperiod of 16 h light and 8 h dark during the whole duration of egg development and a group of eggs was kept in complete darkness. Each light exposed group was kept under different irradiances ranging from high to low: 250, 100 and 10 μmol m^-2^ s^-1^ measured with a SpotOn quantum sensor. Water was changed biweekly with fresh Bat Lake water and weekly for the eggs kept in the dark. The period was decided by measuring ammonia and NH_3_^+^ concentration in the water by using a commercial kit (Freshwater Master Test Kit, Aquarium Pharmaceuticals Inc., Summertown, UK). It is worth noting that light treated egg masses did not present ammonia above 1 PPM (Figure S2).

For sampling of photosynthetic activity, samples were taken out of the temperature-controlled room and kept in an in-house made incubator kept at 15°C in the dark. Measurements of photosynthetic activity and imaging were done at 20°C (given the dimension of the instruments used) and handling lasted approximately 3 minutes.

### 2.3 Progression of embryological and algal development via digital imaging

The stage of embryonic development was tracked for every egg using an in-house developed microscope composed of a SV 305C camera (Svbony, China), a generic zoom lens (Zoom Camera Lens, 25mm Zoom Adjustable Magnification, QANYEGN, Amazon.ca) and a white light LED ring (Tech 1, Dollarama, Montreal, Canada) as source mounted on a retail stand fixed on a lab jack used as Z-axis adjusting stage. Samples were manipulated with a XY stage (Newport Corp, NY, USA) to centre the samples in the field of view. Development was assessed based on observable anatomical features, including the formation of gills, limbs, and tails by using the chart presented by (Harrison 1969). Image acquisition was performed using SharpCap software (Version 4.0; Glover, 2018–2021)

Acquired images were analyzed using FIJI/ImageJ. Images were calibrated with a micrometre slide with 0.01 mm resolution. Image segmentation was used to measure the orthogonal area of the embryo and the algal growth. Algal growth was determined by using the blue channel and segmenting out the area where high absorption of blue light occurred (i.e., dark pixels). Areas were the embryo occurred were not considered in the algal growth analysis. We opted for measuring the pixel intensity in the blue channel of the images because green algae absorb blue light strongly due to the Soret band of chlorophylls *a* and *b* (Zavafer et al. 2023). Because the egg capsule without algal growth is transparent in the visible spectrum, higher concentrations of chlorophyll manifested as dark pixels in the blue region (due to higher light absorption). The size of the areas displaying chlorophyll was used as an indicator of algae colonization.

### 2.3 Algal Growth and Photosynthesis

Chlorophyll *a* fluorescence was measured using two types of instruments depending on the concentration of the algae. When Oophila concentration was low and the egg capsule looked clear, we used a Walz Water PAM (Heinz Walz GmbH, Effeltrich, Germany); when the embryos grew and Oophila concentration increased to the point it was not possible to measure it with the Water PAM, we used an Open-JIP fluorometer (Bates et al. 2019) using a cuvette configuration as published by Bates et al. (2023). Both instruments used blue excitation of 475 nm with a saturating pulse higher than 3000 μmol m^-2^ s^-1^. Measurements of F_O_ (initial level) and F_M_ (maximum level) were used to calculate derivative parameters such as variable fluorescence (F_V_ = F_M_ - F_O_) (Kalaji et al. 2014; Kalaji et al. 2017) and a parameter that links to the net efficiency of photosystem II (F_V_/F_M_) (Sipka et al. 2021).

### 2.6 Analysis of OJIP curves

When samples were measured with the Open-JIP fluorometer we were also able to record the polyphasic rise of the fluorescence commonly referred as OJIP curves. For the analysis of the transient curves, we used the normalized variable fluorescence:

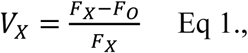

where *V_X_* is the normalized fluorescence, F is the fluorescence intensity at the *X* step, which could be J or I (2 or 20 ms). The J-step represents the inflection point where all QA (primary electron acceptor of Photosystem II) reduction level has been achieved under continuous illumination (Osmond et al. 2017), and I represents when all the plastoquinone (PQ) pool has been reduced (Toth et al. 2007). This biophysical approach correlates with the degree of reduction of Photosystem II (PSII) electron acceptors at the J and I step, which allows for determination of bottlenecks at QA→QB (V_J_) and plastoquinone (PQ) pool/cytochrome b_6_f (V_I_) in the electron transport chain.

Double normalized fluorescence between F_J_ and F_O_ was used to determine the status of the donor side (*e.g,* Mn_4_O_5_Ca cluster) (Strasser 1997) by calculating the so-called K-band (occurring at 300μs after illumination). When two curves are compared, a higher value of the K-band in one curve indicates a less efficient electron donation in PSII (Strasser 1997). To determine the magnitude the of the K-band, the area under (if positive) or above it (if negative) was measured.

### 2.5 Mathematical modeling of the bioenergetics of the holobiont

Holobiont O₂ evolution and consumption rates were predicted under our experimental light conditions by determining the e electron transport rate at different irradiance (as explained in Ralph and Gademann (2005)). Our calculations had the following two assumptions:

- We assume our embryos behaved similarly to those reported by Small et al. (2014) as ETR determinations were carried out during stage 19 to 20. In their work, O₂ consumption by eggs in the dark was 25 μmol egg^-1^ h^-1^ (or 0.58 μmol O_2_ egg^-1^ min^-1^, as calculated here) and under the light (70 μmol m^-2^ s^-1^) net egg oxygen consumption was -10 μmol O_2_ egg^-1^ h^-1^ at 15°C under atmospheric pressure. This means algae evolve ∼35 μmol O_2_ egg^-1^ h^-1^ or 0.58 μmol O_2_ egg^-1^ min^-1^ (calculated by the difference between the two values).
- The light level in Small et al. (2014) is equal or close to photon fluence rate (4π) as explained in Zavafer et al. (2023).

To estimate the maximum photosynthetic rate of O_2_ evolution (Pm), rapid light curves (RLC) were measured in eggs during stage 15 to 19 by using pulse amplitude modulated (PAM) chlorophyll *a* fluorescence. This range of stages was selected because the egg capsule remained transparent which is required to prevent reabsorption of the fluorescence (Serôdio and Campbell 2021). The RLC method (Ralph and Gademann 2005) determines the relative electron transport rate (rETR) pumped by PSII at discreet photon fluence rates (*PFR*) by using the following calculation:

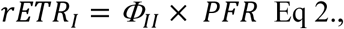

where Φ_II_ is the quantum yield of PSII at the at a given PFR.

Then values of rETR were fit to the Weber model for photosynthesis:

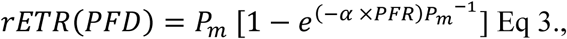

where *α* is coefficient of photoactivation. The values of *rETR* are reported in e^-^ μmol m^-2^ s^-1^, but they are relative to sampled cross section, thus the area of the egg was estimated using a sphere and data is presented as per egg.

The Weber model was used to calculate rates of maximum O_2_ evolution (*rO_2_*) by dividing the theoretical maximum molar ratio *e^-^* to O_2_ which is 4 (which is the stoichiometric value for O_2_ evolution). Because RLC measures the short-term capacity of electron transport (ETR) under quickly increasing light steps, it captures only the immediate, non-steady-state response (Ihnken et al. 2010). For this reason, two scenarios were calculated based on efficiency of the electron transport chain, a pessimistic (60%) and optimistic (80%), both efficiencies within what was observed when compared RLC and steady state photosynthesis in green algae (Flameling and Kromkamp 1998; White and Critchley 1999; Figueroa et al. 2003).

To convert the normalized values of *rO_2_* in absolute numbers, we applied a correcting factor (*C_f_*) based on Small et al. (2014) which is calculated as 0.58/ *nETR*_70_ or electron transport rate at 70 μmol m^-2^ s^-1^ as done below:

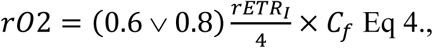

where, 0.6 or 0.8 represents fractional efficiency and 4 is the electron to O_2_ molar ratio. The simulation was done in range of irradiances of 0 to 2000 μmol m^-2^ s^-1^, which is naturally relevant. Calculation was done using a Python 3.12 script and it is available in our Git Hub: [biophotonicsbrock · GitHub](https://github.com/biophotonicsbrock)

### 2.6 Statistical analysis

All the data was analysed using a combination of two software suites: Microsoft Excel, Origin Pro 2023b and the Python Library “Scientific Python”. Specific statistical tests used are disclosed in the legend of every figure.

## 3. Results

### 3.1 Salamander Developmental Rate

To measure developmental rate in the eggs, we assessed the stage of development between groups, and the results are displayed in Figure 1A, where the mean, maximum and minimum stage are displayed for each treatment. Embryonic developmental speed coincided with the presence of light, where the low light treated eggs displayed slower development (10 stages behind high light at experimental day 25) and hatching than the light treated samples (taking up to 45 days to hatch). Higher and medium light treated embryos started to hatch only after 33 days of the experiment. (Figure 1B). Dark treated eggs were the slowest in development displaying a delay of 1 week against the high light eggs and a smaller size (Figure 2). A comparison between the embryos at the same stage of development (stage 40 during day 43) between low light and dark treated samples shows the latter group had a smaller size (see Figure 3).

**Figure 1.**
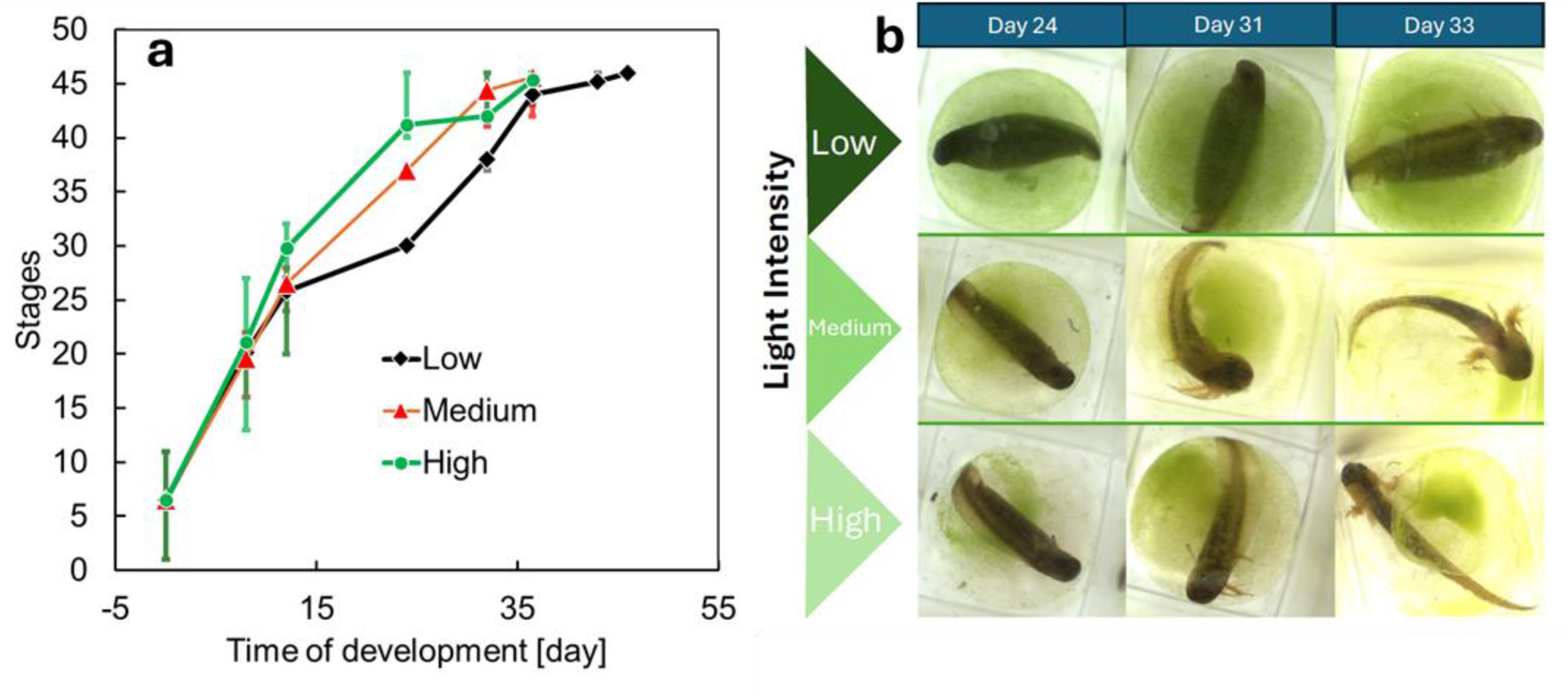
Embryos develop faster in high light, but at low light there is more algal growth. **(a)** Comparison of micrographs taken on the same day showing differences in embryo development, each datapoint correspond to the average of > 7 eggs and error bars represent minimum and maximum stage of development. **(b)** Comparison in the progression of development during the experiment for the three light intensities. Ordinal data of stages was analyzed using an ordinal regression (cumulative link models, via statmodels Python, output in Table S1, day (coef = 0.71, p < 0.001), group ML (coef = 2.86, p < 0.001) and coef = 2.33, p = 0.001), * means both ML and HL groups develop faster than LL, and all groups progress with time.

**Figure 2.**
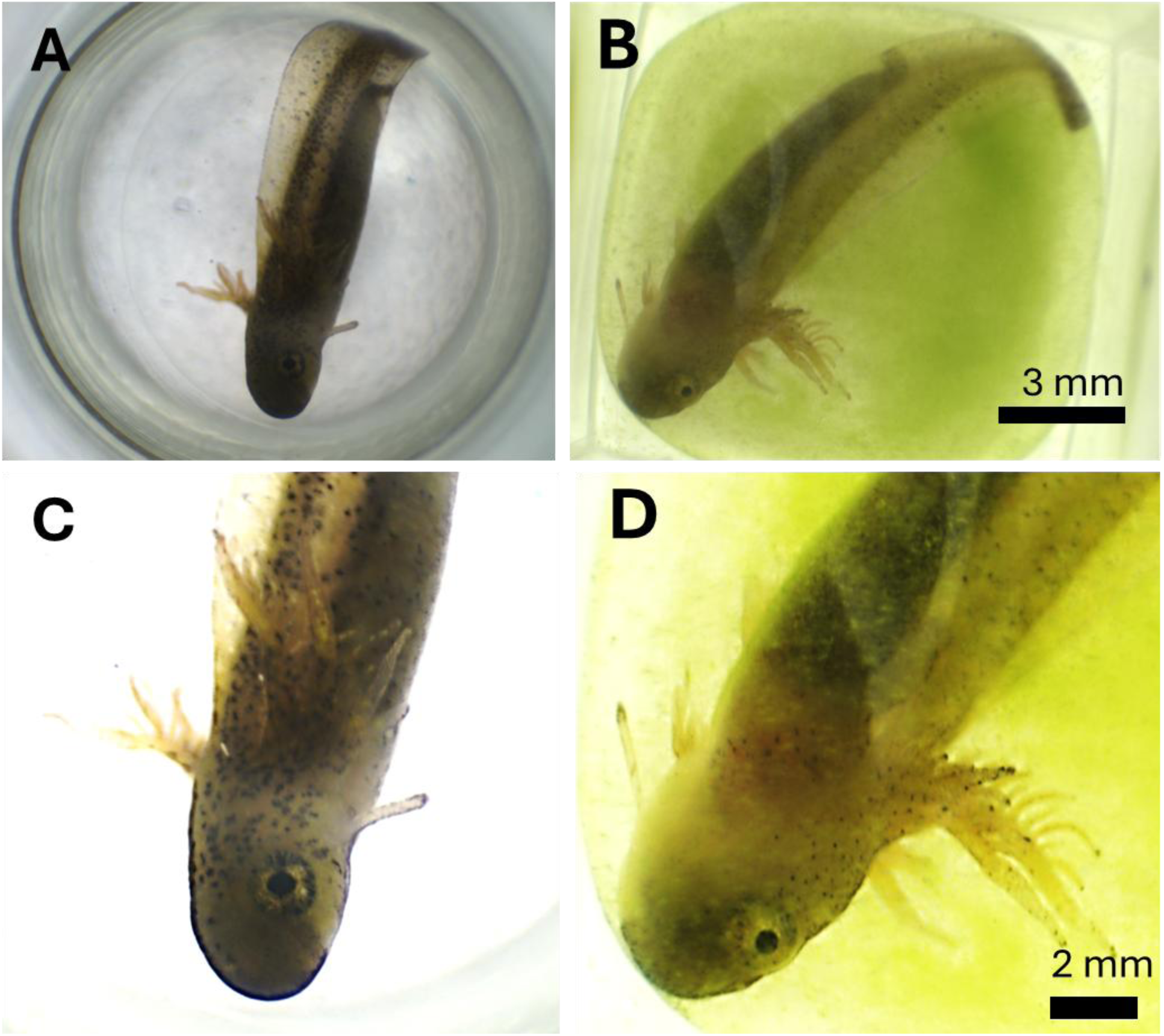
Comparison of the development of two representative specimens of dark-(left column) versus low light-treated (right column) embryo at developmental stage 40. Whole eggs of dark-treated **(a)** and low light-treated **(b)**. **(c-d)** Zoom-in of the upper portion of embryos to illustrate size difference, for these images an enhancement of contrast and sharpness was applied to compensate against light scattering by the algae. Dark-treated (c) and low light-treated (d) embryo.

**Figure 3:**
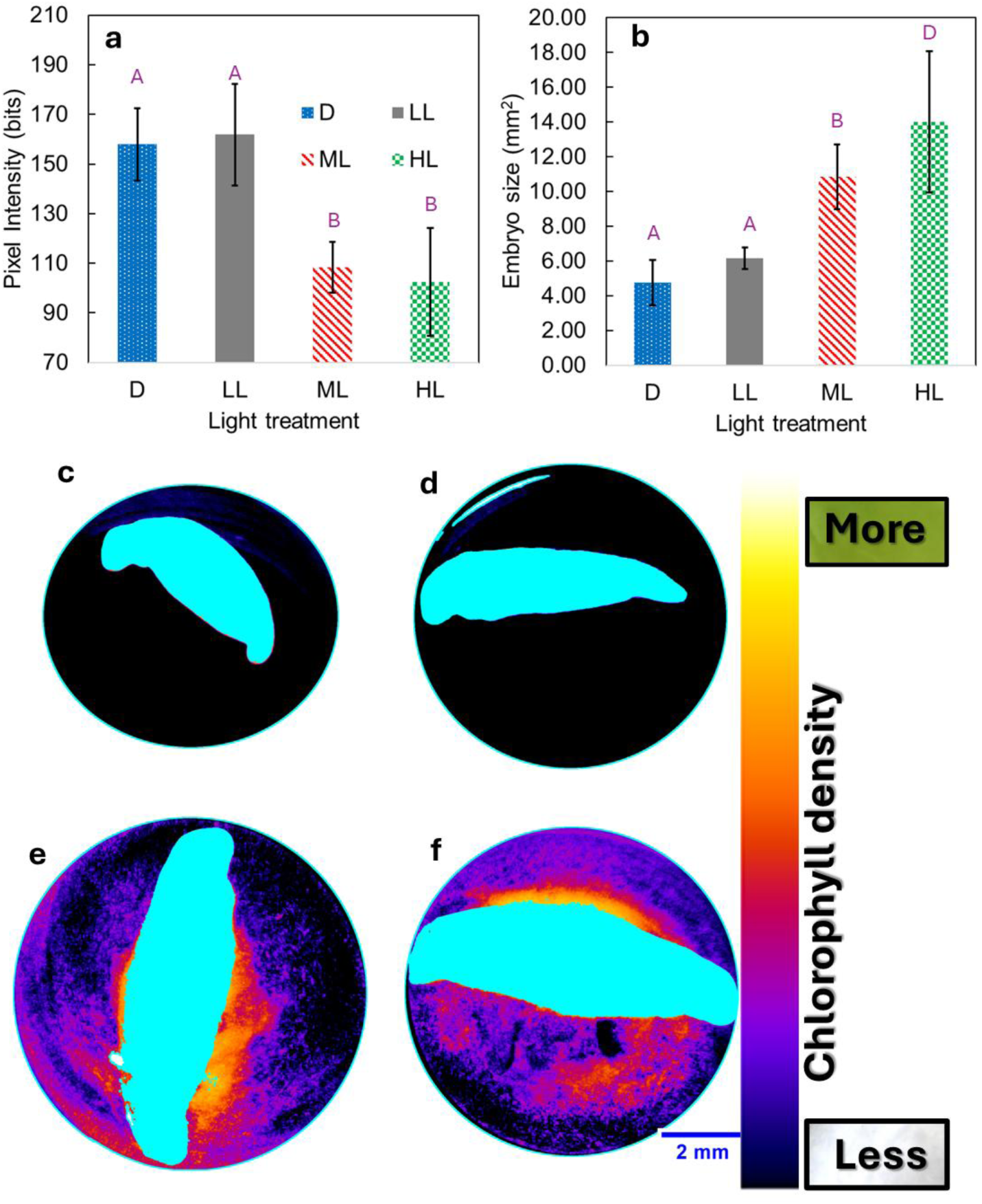
Growth of the holobiont measured by digital micrography analysis. Data corresponds to experimental day 20. **(a)** Light intensity in the blue channel of RGB images in the non embryo region, where the darker a group is where more blue light chlorophyll absorbs. **(b)** Embryo size determined by measuring the area in each micrograph using the green channel exclusively. Experimental light conditions: dark (D), low light (LL), medium light (ML) and high light (HL). Each data point corresponds to the average ± standard deviation of n > 11 per group. Letter-group scheme for treatments sharing a letter is NOT significantly different, and vice versa. A one-way ANOVA was implemented with Tukey HSD post-hoc test with α = 0.05. **(c-f)** Segmentation analysis of characteristic eggs displaying embryos (in cyan) and algal spreading by in the egg in false color. Dark (C), low- (D), medium-(E) and high-light (F). A false color look-up table was implemented where “hot” represents a higher chlorophyll density in arbitrary units. Rectangular frames displaying “More” and “Less” show the actual color of the images analyzed.

### 3.2. Algal Growth Density

We could not detect algal growth in dark treated eggs using fluorometry and microscopy even in later days of the experiment (day 43, Figure 2A). In all light treated samples, algae colonization was observed (Figure 1A and 2B-D).

By using the images captured in days 17 and 18, we were able to determine the level of algal growth in the three light treatments, where growth within the egg capsules showed an inverse relationship with irradiance levels at day 20 (Figure 3A). Low light exposed eggs had low algae density (Figure 3A) and were not statistically significant when compared with dark treated eggs. In some low light treated eggs, algae were detectable under the microscope, which was confirmed in the photochemistry measurements (Figure 4A). Medium and high light treated eggs displayed clear algal colonization under the microscope which was significantly higher than low and dark treated eggs. It is worth noting that the average area of the embryo increased as the light levels increased.

**Figure 4:**
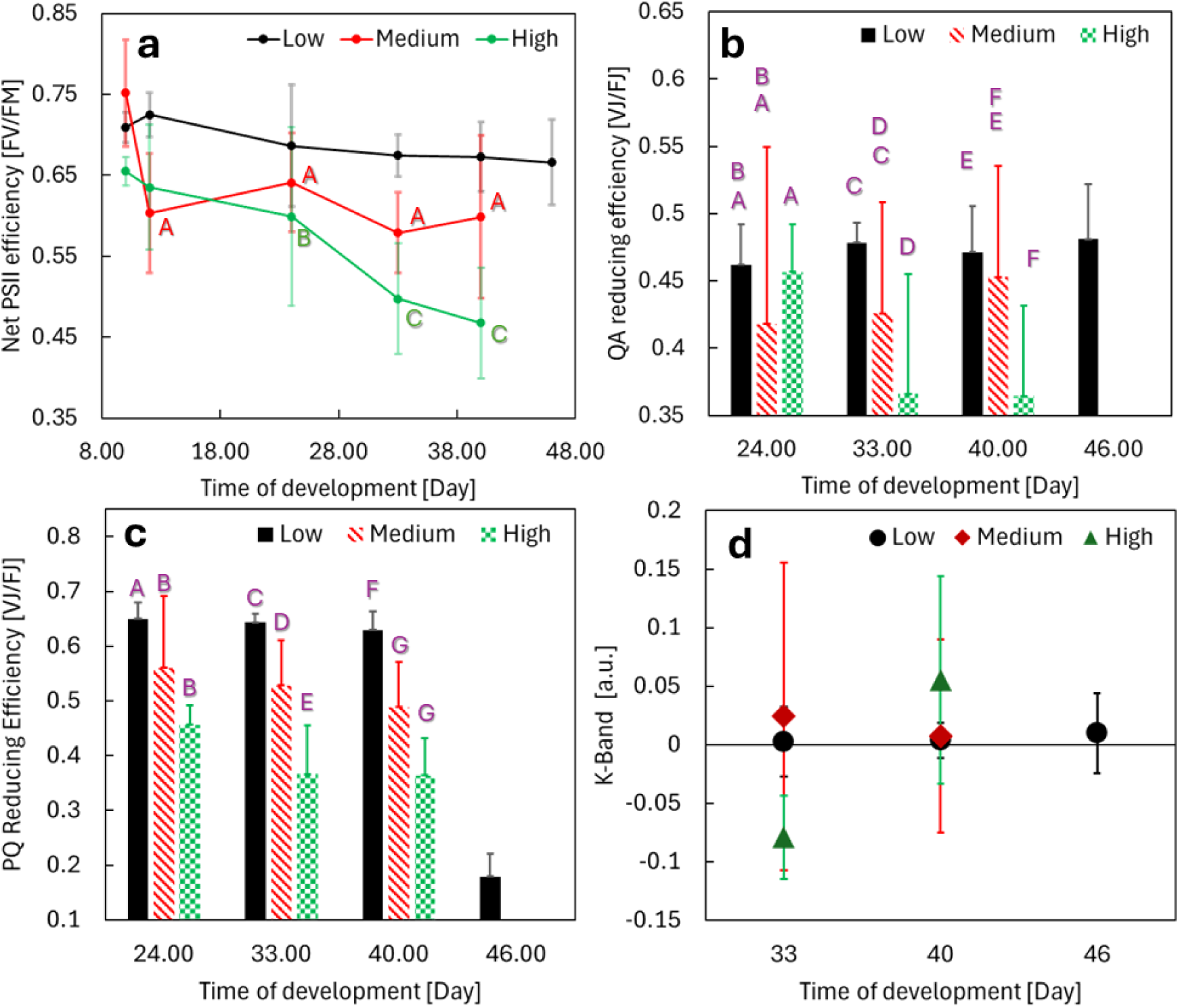
Biophysical parameters of PSII performance estimated from chlorophyll *a* fluorescence. **(a)** Time course of F_V_/F_M_, representing PSII net quantum efficiency. No significant difference in time was observed for low light. First, statistical comparison between treatments using the first time point was done using one-way ANOVA was implemented with Tukey HSD post-hoc test with α = 0.05. **(b)** Time course of change of V_J_/F_J_, representing Q_A_ reducing efficiency. For b, HL was used as baseline, and effect was found after day 40 between LL and ML (p < 0.05). **(c)** Time course of change of V_I_/F_I_, representing PQ pool reducing efficiency. For c, HL was used as baseline, and effect was found after day between LL and ML (p < 0.05). **(d)** Integrated area of the K-band at three different time points, as a point of reference the OJIP curves during day 24 were used. Statistical analysis of panels a-c was done using a generalized estimating equations (GEE) regression. Each data point is the average ± standard deviation of n > 7. No significance was found among data point.

No widespread algae colonization was found in dark and low light treated eggs (Figure 3CD) but was abundantly found in medium and high light treated eggs (Figure 3EF). We can exclude localized thermal effects from the LED lights influencing these results, as we confirmed our light system lights did not increase the temperature of the samples.

### 3.3 Net and Partial Photochemical Efficiency of Light Reactions

Oophila photosynthetic performance was monitored *in situ* by measuring the changes in F_V_/F_M_, a biophysical parameter that correlates with the net efficiency in how PSII pumps electrons through the light reactions of photosynthesis (Figure 4A). A GEE analysis revealed no significant differences among treatments at the first time point. Value decreased significantly over time (p < 0.001). Significant interaction in the light treatment and the time indicated that low-light (p < 0.001) and medium-light (p = 0.012) declined less steeply than high-light, suggesting that these treatments mitigated the temporal decrease in value. No photosynthetic signals were observed in dark treated eggs even under high gain in the Water PAM fluorometer.

Decay of F_V_/F_M_ could be caused by strong alterations in the function of PSII, either by down regulation of photosynthesis or by actual destructive stress, and this can be inferred from the chlorophyll emission kinetics. Thus, we used normalized variable fluorescence (V_x_) analysis to find the causes behind the net loss of photosynthetic efficiency. Phases O-J and J-I correlate with different inflection points of the OJIP curve and link with steps of photosynthetic electron transport chain. Polyphasic rise of the chlorophyll fluorescence was recorded from day 24 and the partial ratios (V_J_/F_J_ and V_I_/F_I_) in Figure 4BC. For V_J_/F_J_ (Figure 4B) a GEE analysis showed no treatment differences at Day 24. Across all groups of Figure 4b, values decreased over time. However, low-light and medium-light treatments exhibited significant positive interactions at Days 32 and 40, indicating that these groups maintained or increased values relative to high-light, which continued to decline, indicating that reduction of QA by PSII activity was stable. For, V_I_/F_I_ (Figure 4C) a GEE analysis revealed that medium-light treatment started significantly lower than high-light at baseline. Across all groups, values declined over time. However, low-light and especially medium-light showed positive treatment and time interactions, indicating that these groups declined less steeply and medium-light improved relative to high-light as time progressed. This means that with low-light conditions, medium-light eggs exhibited reduced PQ-reducing efficiency beginning at day 24, while high-light eggs showed lower V_I_/F_I_ values as early as 24 h (Figure 4C). With respect to temporal effects, V_I_/F_I_ in low-light eggs remained unchanged until day 48 (Figure 4C). Both medium- and high-light treatments showed no statistical differences between days 33 and 40 beyond decrease after day 24 (Figure 4C).

A decline in QA-reducing efficiency as a function of increase of light could be ceased by photoinhibition (Osmond and Grace 1995). PSII photoinhibition may be caused by damage to the water-splitting site (PSII donor side) (Hakala et al. 2005) or the acceptor side limitations (QA reducing side) (Vass and Cser 2009). To evaluate donor-side performance, K-band analysis was conducted (Figure 4D). Although a positive K-band was detected across all treatments at day 40, values did not differ significantly from those recorded at day 24 within each treatment. Although medium- and high-light conditions produced an average positive K-band when compared to low light the differences were not statistically significant. All this indicates that impairment of the Mn_4_O_5_Ca cluster can be discarded as the main driver of the observed decay of photosynthesis activity. Because light induced damage of PSII (chronic photoinhibition) commonly leads to the formation of K-band (Zavafer 2021), down regulatory photoinhibition (dynamic photoinhibition) is the likely explanation of the observed decay of PSII activity (Osmond and Grace 1995).

Dynamic photoinhibition often changes the shape of the OJIP curve due to modulation of the electron transport chain. Thus, we examined the shape of the whole OJIP curves to determine if other effects could be detected (Figure 5). We observed an overall increase in the OJ phase as light intensity increase, and in parallel a decrease in the IP phase. Temporal effects in shape were minor between low and medium light treated eggs (Figure 5bc); however, OJIP of the high light treatment showed an absence of the JI phase (Figure 5d), which is explained by diminished electron transport.

**Figure 5:**
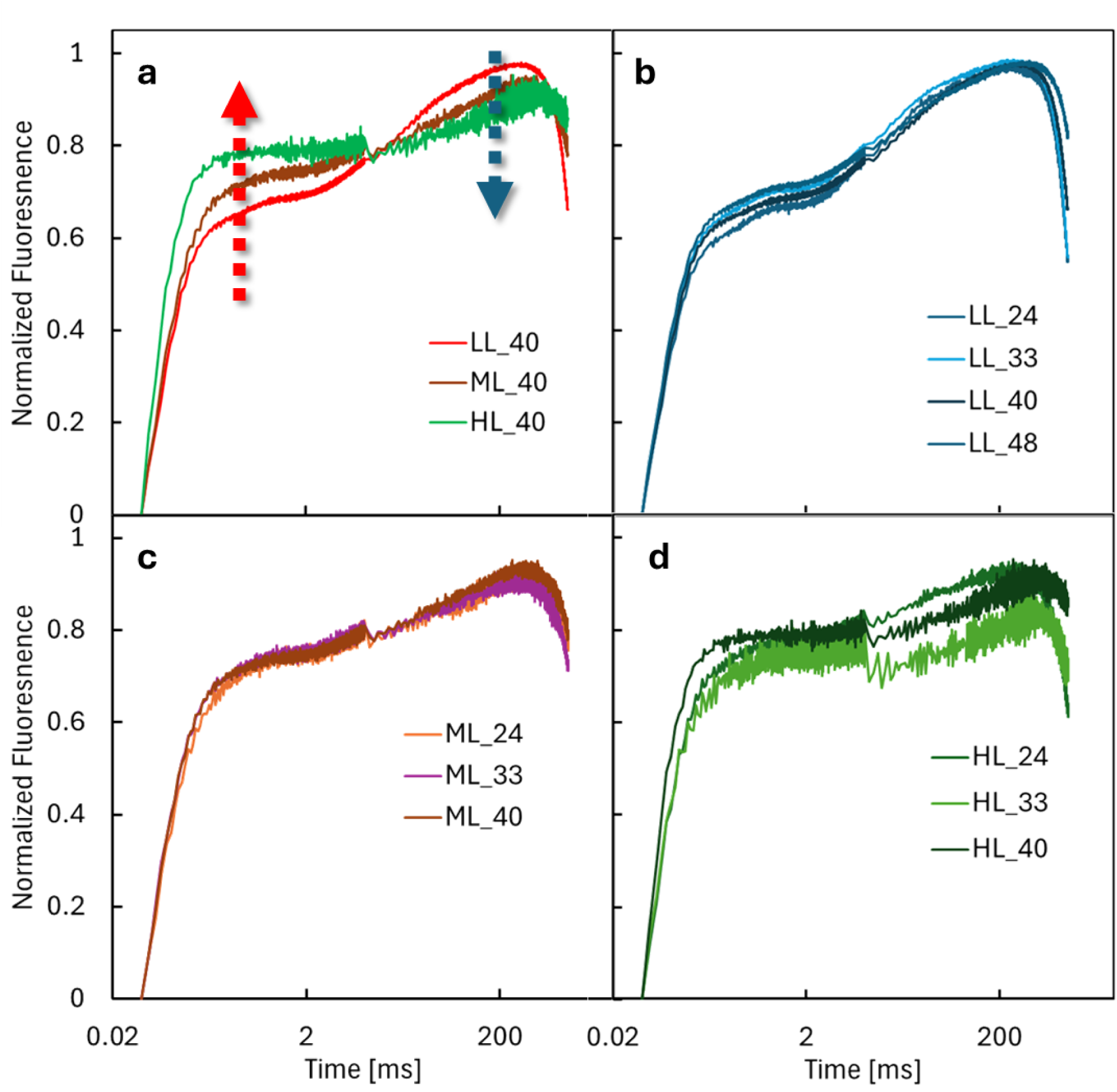
Average OJIP curves of different time points and treatments. **(a)** Comparison of the OJIP curve at 40 days between the three light treatments. Red arrows highlight an overall increase of the OJ phase, while blue arrow highlights loss of the IP phase as irradiance increased. **(b)** Time series of low light (LL) treated eggs. **(c)** Time series of medium light (ML) treated eggs. **(d)** Time series of high light (HL) treated eggs. OJIP curves were double normalized (F_O_ = 0, F_M_ = 1), Each data point is the average ± standard deviation of n > 7.

### 3.4 Predictive analysis of O_2_ evolution and consumption in the holobiont

To determine if the O_2_ produced by the algae is enough to support the needs of the embryo, a simulation to estimate O_2_ produced and consumed by the holobiont was performed (Figure 6). To do this, experimental RLC determinations were carried out, and the result of the electron transport rate (ETR) is presented in Figure 6A. The experimental points adjusted well to the Weber model (Figure 5A) yielding a Pm of 193.25 ± 12.74 μmol e^-^ m^-2^ s^-1^, α of 0.09 ± 0.002 with a r^2^ 0.998. These values were normalized to the calculated Pm and corrected according to Eq 4. The simulation predicts that O_2_ evolution by the algae at high light would exceed the basal O_2_ consumption by three times of the embryo at 60 and 80% efficiency. At medium light, O_2_ production would exceed the basal O_2_ consumption by 20% to support average O_2_ consumption by the embryo, while low light was far below the embryo O_2_ requirements (Figure 5b).

**Figure 6:**
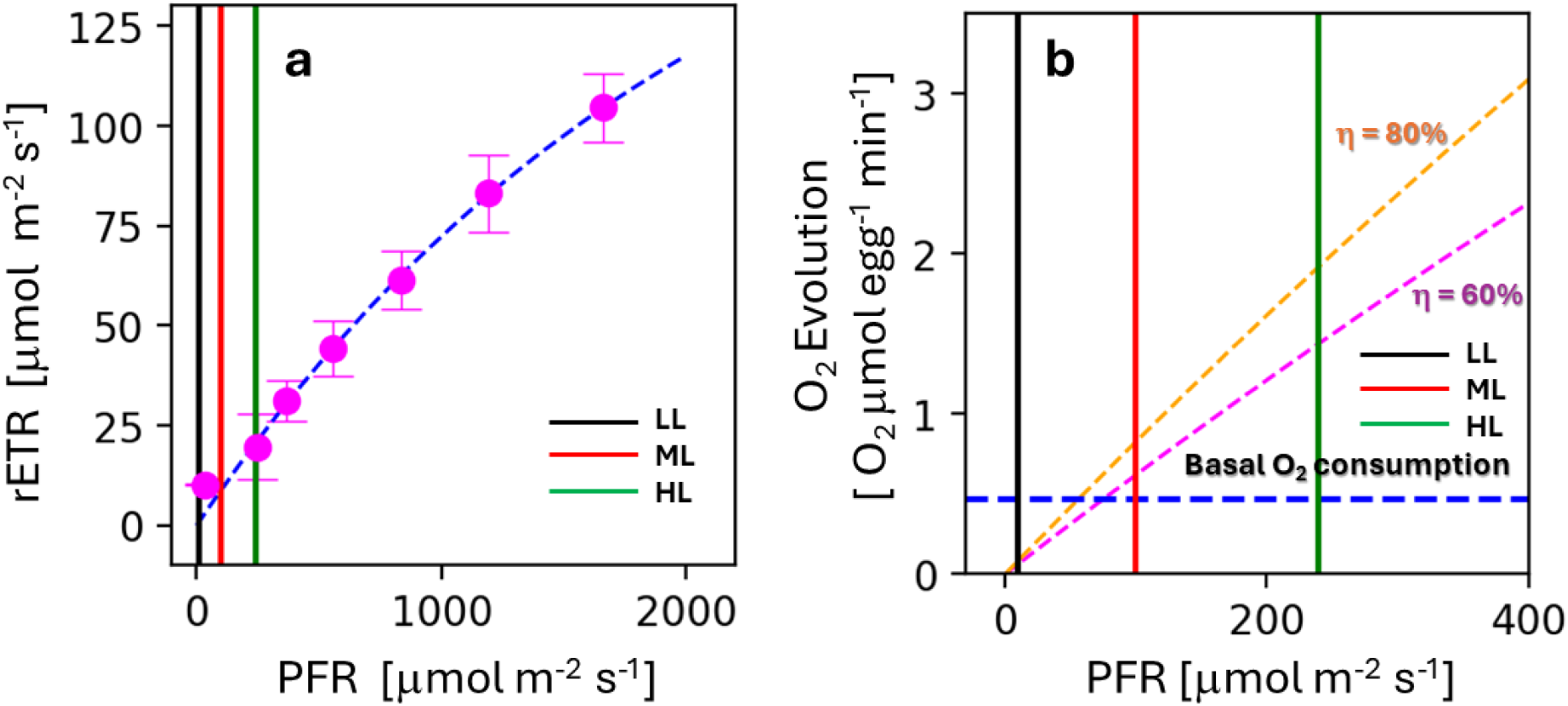
Photosynthetic capacity of eggs (stages 19 to 20) and scenarios of electron transport of the holobiont. **(a)** Experimental determinations of relative electron transport rate responses as a function of photon fluence rate, in magenta average experimental determinations of 3 eggs from different egg masses, blue dotted line represents Weber fitting. **(b)** Simulation of O_2_ evolution as a function of PFR, two scenarios of O_2_ evolution are considered based on net efficiency (η), 80% in dotted orange and 60% in magenta, basal O2 consumption by the embryo is represented by the blue line. In both panels the vertical lines represent our experimental light conditions: low light (LL), medium light (ML) and high light (HL).

## 4. Discussion

In this work, we have investigated the effect of light dose on the development of embryos and growth of algae, in the context of their symbiotic relationship. Our experiments evaluated two explicative paradigms, direct mutualism and algal stress (a type of amensalism). Literature supporting the shift to fermentative metabolism (here referred as the algal stress paradigm) alludes to the idea that Oophila algae inside salamander embryos shift away from photosynthetic due to the low-oxygen and low-light intracellular environment (Burns et al. 2017). Here, we observed that light dose influences Oophila grow in the egg capsules of spotted salamanders. If Oophila is not mainly growing photosynthetically, as the stress paradigm proposes (Burns et al. 2020; Burns et al. 2017), different light treatments would have minimal impact on algal growth and performance. Although, Burns et al. (2020) showed that salamander embryos themselves fix significant amounts of inorganic carbon, making it difficult to detect any substantial transfer of photosynthate from Oophila to the host, which strictly speaking does not negate photosynthesis.

Our experiments strongly point light is a major driver of growth for Oophila, which is supported by other works (Hutchison and Hammen 1958; Small et al. 2014; Small and Bishop 2020). We did not observed signs of fermentation like in other works (Burns et al. 2020; Burns et al. 2017), as we should have seen green algae in the dark treated eggs, but ferementation was not measured directly. However, Chlamidomonales shall express chlorophyll even in the absence of photosynthesis (Duanmu et al. 2013) when growing via fermentative pathways.

Through *in situ* and *in vivo* monitoring of algal photosynthesis, we found an inverse relationship between conditions that favour host developmental rate and those that optimise PSII performance. We found algal abundance, developmental rate and size of the embryos coincided. All which agree with mutualistic endosymbiosis. Specifically, high light exposure accelerated developmental speed of embryos and algae.

Despite the parallel changes in growth of algae and embryo, PSII activity was the lowest in high light treated eggs. This result can be counter intuitive under the mutualistic paradigm, as accelerated growth of algae should be caused by higher rates of O_2_ evolution by the algae, which should occur when PSII is more active (Hutchison and Hammen 1958; Small et al. 2014). However, in green algae low values of F_V_/F_M_ could indicate lower rates of PSII activity due dynamic photoinhibition (Osmond and Grace 1995; Han et al. 2000). Dynamic photoinhibition occurs when the net excitation exceeds the current energetic needs of the photosynthetic organism and the activity per PSII unit is modulated downwards (Han et al. 2000). The other type of photoinhibition, so-called chronic (Osmond and Grace 1995), involves damage of PSII (Zavafer et al. 2015) and can be discarded as an explanation because no clear K-band was detected, an indicator typically associated with PSII photodamage to the Mn₄O₅Ca cluster (Iermak et al. 2020).

Micrographs show abundant algae in the eggs and that would lead to higher bulk concentrations of O_2_ in the holobiont, although the photosynthetic activity per PSII protein complex is decreased. In fact, this interpretation is supported by the two works of Small et al., first net O_2_ production by the holobiont increases as the developmental stages advance (Small et al. 2014); and, by the result that algal growth and chlorophyll concentration decreases under 24 h photoperiods, as long photoperiods often lead to photoinhibition in algae (Small and Bishop 2020). In the scenario of dynamic photoinhibition, the lowest light condition presents the highest PSII activity, as PSII protein complex operate below light saturation. Based on our simulation of O_2_ evolution, rates of electron transport cannot sustain a production of O_2_ enough to support embryonic development by themselves at low light levels, explaining the slower development of embryos.

Previous research supporting fermentation and the stress paradigm have been done using irradiances below that which would support the basal O_2_ consumption by the embryo (Burns et al. 2017; Burns et al. 2020) and can explain why algae expressed fermentation genes. Our results show the relevance of understanding the algal photobiology, as Oophila performance will have an impact on the symbiotic relation between the two species. If the experiments are carried out under conditions where the algae growth is reduced (such as low and no-light conditions), the net symbiotic benefits may be nullified or even reverted into an amensalism.

The interpretation above does not negate the stress paradigm, but does not support it as the sole driver of the symbiosis. It is worth mentioning that medium and high irradiances used here likely exceed the light levels the holobiont would naturally encounter in some ponds (Earl et al. 2011; Rowland et al. 2017). Spotted salamanders typically lay their eggs at depths of roughly 300 mm (Kern et al. 2013), where irradiance is substantially lower. It remains plausible that Oophila experiences physiological stress under certain environmental conditions and, in those instances, expresses the gene sets reported by Burns et al. (2017). Gene expression is likely stage and time of day dependent.

In nature, embryos develop under markedly hypoxic conditions (Valls and Mills 2007). Eggs are typically laid at depths where atmospheric O_2_ exchange is minimal (Kern et al. 2013), and the surrounding protein-rich jelly matrix further restricts diffusion (Burns et al. 2017; Burns et al. 2020). In contrast, our experimental eggs were maintained at a depth of only 15 mm and removed from the jelly, allowing the atmosphere to replenish both CO₂ for photosynthesis and O₂ for embryonic respiration. Nevertheless, atmospheric oxygenation alone cannot account for the positive effects of light on embryo development (Small and Bishop 2020); these observations are more plausibly explained by additional O₂ supplied through algal photosynthesis.

Some limitations in our experimental design and in the interpretation of our results must be acknowledged. Because the fluorometric approach used here captures the average performance of the algal population within the surface of each egg, it is also possible that, algae deeper in the egg and tissue experience other physiological conditions. A further limitation is that all biophysical assessments relied exclusively on fluorometric techniques. While the non-invasive nature of chlorophyll fluorescence enables continuous monitoring of the holobiont, the method reports only on PSII photochemistry and does not directly quantify pigments, proteins, or metabolites (Kalaji et al. 2017; Kalaji et al. 2014). Even gas-exchange measurements would not fully resolve this limitation, as O₂ production by the algae and O₂ consumption by the embryo cannot be disentangled without the use of stable isotopes. Fluorescence signals therefore cannot replace biochemical assays, although they do provide valuable, non-destructive insight into the *in vivo* bioenergetics of the holobiont. Stable-isotope approaches coupled with membrane-inlet mass spectrometry, as commonly applied in plant and coral (Tansik et al. 2017) research, could in principle separate respiratory and photosynthetic fluxes. However, these methods operate at substantially lower throughput, which would limit statistical power and compromise the scope of the study. Given these constraints, chlorophyll fluorescence remains an appropriate and informative tool for assessing *in vivo* photosynthetic performance in holobiont.

Despite the limitations of our work, our data supports the idea that depending on the circumstances both paradigms may be true. An argument to support mutualism could be that spotted salamanders have a wide distribution in North America and can be considered a highly successful organism, having adapted to many ecosystems and stressors. Because a wide distribution represents high fitness, the occurrence of endosymbiosis in all populations may be part of this success. This idea is clearly influenced by other mutualistic symbiosis of microalgae and animals, e.g., corals (Mohamed et al. 2016).

## 5. Conclusions

Our work contradicts certain fundamentals of the stress and mutualistic paradigm. Our experimental observation that algae failed to grow in dark-exposed eggs confirms the biological necessity of light for algal viability. Since algae clearly are growing photosynthetically, the premise of growth as fermentation as the main algal driver is refuted. Mutualism on the other hand seems to be circumstantial and a persistent ambiguity remains regarding the exact benefit derived by the algae.

## CreDit statement

Conceptualization: AZ, GT; Methodology: AZ; Software: AZ, HB; Validation: LP, AZ; Formal analysis: AZ; Investigation: LP, AZ, HB, WEC, GT; Resources: PM, AZ, HB, GT; Data Curation: AZ, LP; Writing - Original Draft: LP, AZ; Writing - Review & Editing: LP, PM, GT; Visualization: LP, AZ; Supervision: AZ; Project administration: AZ; Funding acquisition: AZ, GT.

## Acknowledgement

LP and AZ thank Dr. Jae Jung for the support in the preparations for the field work in Algonquin Park. AZ and GT would like to thank the staff of the Algonquin Park Research Sation for their continuous and selfless support.

