## Supplementary Files for "MUTALISM BETWEEN *Ambystoma maculatum* AND *Oophila amblystomatis* IS DEPENDENT ON LIGHT INTENISTY DURING PHOTOSYNTHETIC GROWTH"

SUPPEMENTARY DATA

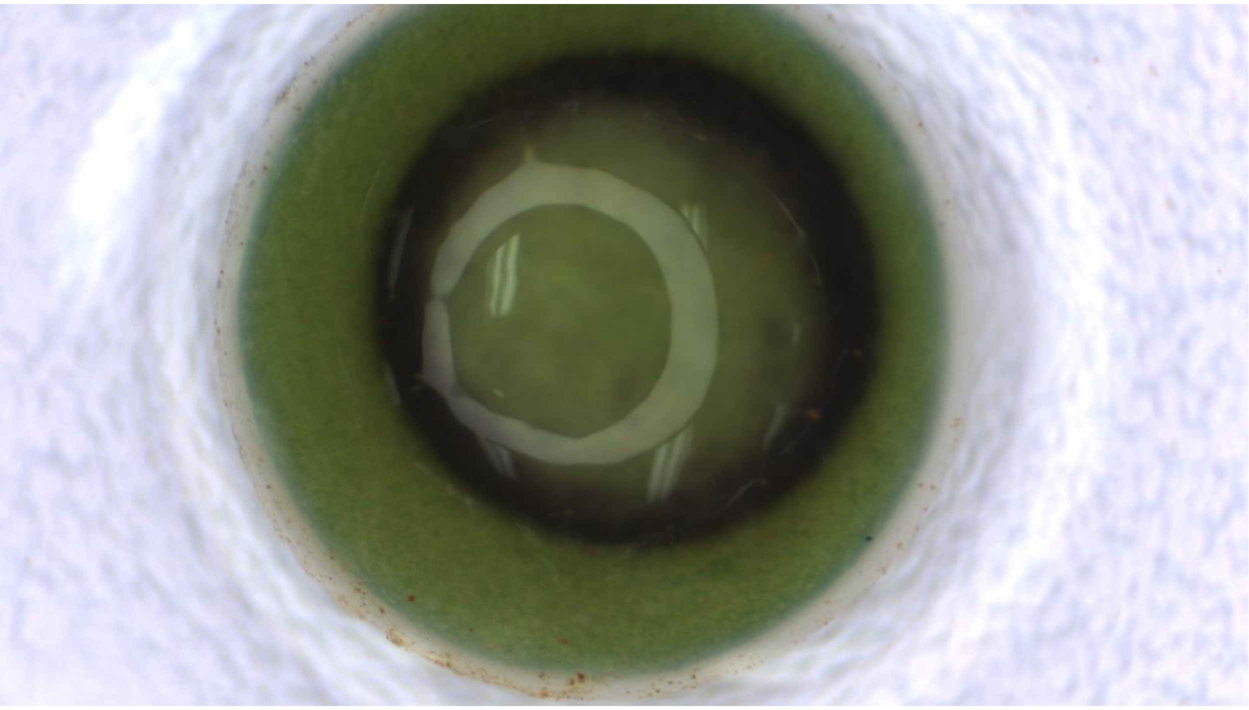

**Figure S1:** A salamander egg where the embryo was not viable and a saprophyte fungus developed in their interior. This image was taken on experimental day 16. It is worth noting that the algal growth was more abundant than those observed for viable eggs.

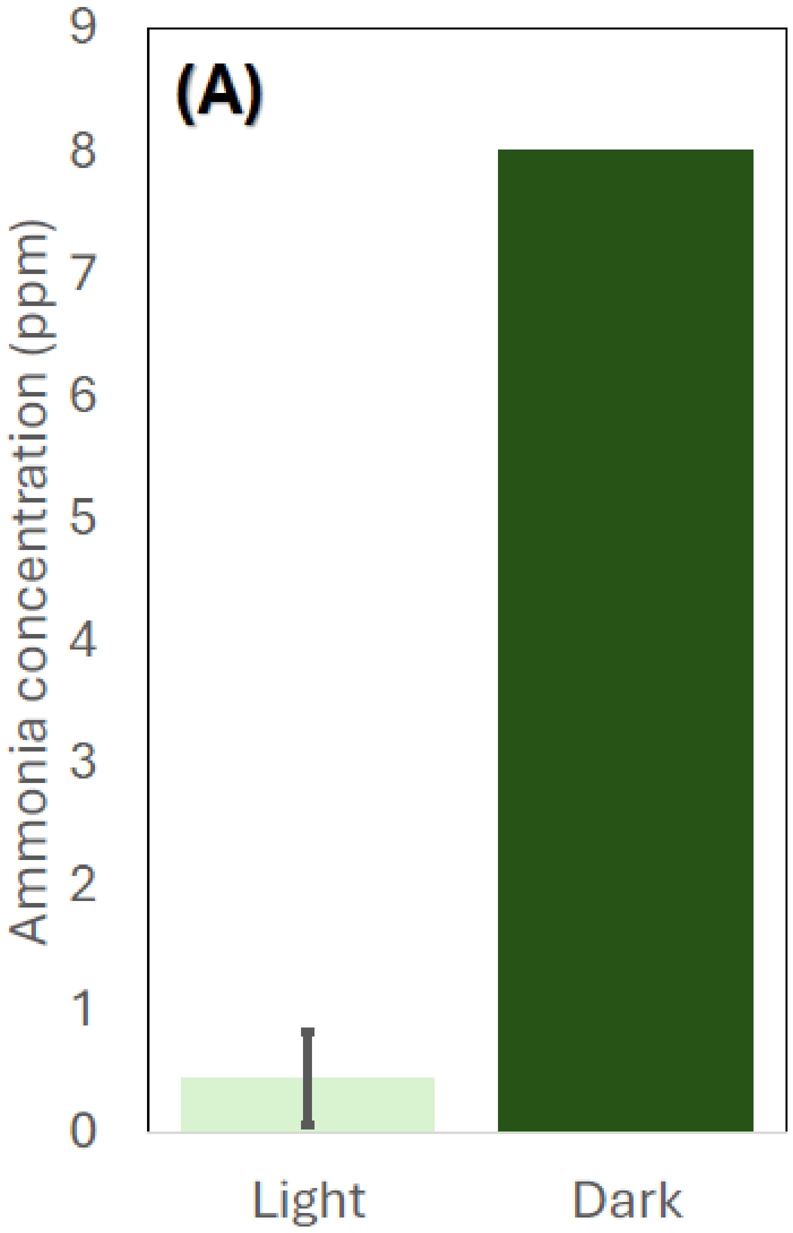


**Figure S2:** Measurement of  concentration NH4 in the water where egg masses were kept.

Table S1. Statistical analysis Figure 1a. Python Output

```
                               GEE Regression Results
===================================================================================
Dep. Variable:                          Value   No. Observations:              177
Model:                                    GEE   No. clusters:                   43
Method:                           Generalized   Min. cluster size:               1
                        Estimating Equations   Max. cluster size:               5
Family:                              Gaussian   Mean cluster size:             4.1
Dependence structure:            Independence   Num. iterations:                 2
Date:                        Thu, 24 Sep 2026   Scale:                       0.004
Covariance type:                       robust   Time:                     01:13:29
=======================================================================================
                         coef    std err          z      P>|z|      [0.025      0.975]
---------------------------------------------------------------------------------------
Intercept              0.7557      0.026     29.435      0.000       0.705       0.806
Treatment[T.LL]       -0.0195      0.029     -0.666      0.506      -0.077       0.038
Treatment[T.ML]       -0.0204      0.030     -0.687      0.492      -0.079       0.038
Day                   -0.0075      0.001     -6.295      0.000      -0.010      -0.005
Treatment[T.LL]:Day    0.0057      0.001      4.475      0.000       0.003       0.008
Treatment[T.ML]:Day    0.0032      0.001      2.505      0.012       0.001       0.006
==============================================================================
Skew:                        -1.2695   Kurtosis:                         2.7858
Centered skew:               -0.6422   Centered kurtosis:                1.7469
==============================================================================
```

Table S2. Statistical analysis Figure 1b. Python Output

```
                             OrderedModel Results
==============================================================================
Dep. Variable:                  stage   Log-Likelihood:                -95.262
Model:                   OrderedModel   AIC:                             212.5
Method:            Maximum Likelihood   BIC:                             237.1
Date:                Wed, 23 Sep 2026
Time:                        00:59:26
No. Observations:                  69
Df Residuals:                      58
Df Model:                           3
==============================================================================
                 coef    std err          z      P>|z|      [0.025      0.975]
------------------------------------------------------------------------------
day            0.7122      0.109      6.533      0.000       0.499       0.926
group_ML       2.8584      0.616      4.644      0.000       1.652       4.065
group_HL       2.3299      0.715      3.257      0.001       0.928       3.732
8/30          17.0222      2.764      6.158      0.000      11.605      22.440
30/37          0.6209      0.274      2.268      0.023       0.084       1.157
37/38          0.9395      0.264      3.553      0.000       0.421       1.458
38/40          0.5022      0.293      1.716      0.086      -0.071       1.076
40/41         -0.5196      0.558     -0.931      0.352      -1.613       0.574
41/44         -1.4079      0.978     -1.440      0.150      -3.324       0.508
44/46          0.7674      0.229      3.349      0.001       0.318       1.217
46/47          0.4019      0.364      1.103      0.270      -0.312       1.116
==============================================================================
```

Table S3. GEE regression for $V_{J}/_{J}$ data Figure 1c.Python Output

```
                               GEE Regression Results
===================================================================================
Dep. Variable:                          Value   No. Observations:                101
Model:                                    GEE   No. clusters:                     38
Method:                           Generalized   Min. cluster size:                 1
                         Estimating Equations   Max. cluster size:                 3
Family:                              Gaussian   Mean cluster size:               2.7
Dependence structure:            Independence   Num. iterations:                   2
Date:                        Wed, 23 Sep 2026   Scale:                         0.003
Covariance type:                       robust   Time:                       16:37:24
=============================================================================================
                                coef    std err          z      P>|z|      [0.025      0.975]
---------------------------------------------------------------------------------------------
Intercept                     0.4565      0.011     42.823      0.000       0.436       0.477
Treatment[T.LL]               0.0067      0.013      0.520      0.603      -0.019       0.032
Treatment[T.ML]              -0.0065      0.019     -0.351      0.726      -0.043       0.030
Day[T.32]                    -0.0905      0.029     -3.097      0.002      -0.148      -0.033
Day[T.40]                    -0.0923      0.032     -2.870      0.004      -0.155      -0.029
Treatment[T.LL]:Day[T.32]     0.1050      0.031      3.424      0.001       0.045       0.165
Treatment[T.ML]:Day[T.32]     0.0663      0.037      1.803      0.071      -0.006       0.138
Treatment[T.LL]:Day[T.40]     0.0998      0.035      2.831      0.005       0.031       0.169
Treatment[T.ML]:Day[T.40]     0.1307      0.037      3.567      0.000       0.059       0.203
==============================================================================
Skew:                        -0.1359   Kurtosis:                         0.7722
Centered skew:               -0.4255   Centered kurtosis:                0.2087
==============================================================================
```

Table S4. GEE regression for VJ/J data.

GEE Regression Results

| | | | |
|---|---|---|---|
| Dep. Variable: | Value | No. Observations: | 101 |
| Model: | GEE | No. clusters: | 38 |
| Method: | Generalized<br>Estimating Equations | Min. cluster size:<br>Max. cluster size: | 1<br>3 |
| Family: | Gaussian | Mean cluster size: | 2.7 |
| Dependence structure: | Independence | Num. iterations: | 2 |
| Date: | Wed, 23 Sep 2026 | Scale: | 0.010 |
| Covariance type: | robust | Time: | 17:54:44 |

| | coef | std err | z | P>\|z\| | [0.025 | 0.975] |
|---|---|---|---|---|---|---|
| Intercept | 0.9607 | 0.210 | 4.583 | 0.000 | 0.550 | 1.372 |
| Treatment[T.LL] | -0.2748 | 0.220 | -1.249 | 0.212 | -0.706 | 0.156 |
| Treatment[T.ML] | -0.7801 | 0.214 | -3.648 | 0.000 | -1.199 | -0.361 |
| Day | -0.0168 | 0.008 | -2.181 | 0.029 | -0.032 | -0.002 |
| Treatment[T.LL]:Day | 0.0155 | 0.008 | 1.944 | 0.052 | -0.000 | 0.031 |
| Treatment[T.ML]:Day | 0.0309 | 0.008 | 3.939 | 0.000 | 0.016 | 0.046 |

| | | | |
|---|---|---|---|
| Skew: | 1.5839 | Kurtosis: | 12.8333 |
| Centered skew: | 0.3011 | Centered kurtosis: | 4.6528 |
